# Interspecies transcriptome analyses identify genes that control the development and evolution of limb skeletal proportion

**DOI:** 10.1101/754002

**Authors:** Aditya Saxena, Virag Sharma, Stanley J. Neufeld, Mai P. Tran, Haydee L. Gutierrez, Joel M. Erberich, Amanda Birmingham, John Cobb, Michael Hiller, Kimberly L. Cooper

## Abstract

Despite the great diversity of vertebrate limb proportion and our deep understanding of the genetic mechanisms that drive skeletal elongation, little is known about how individual bones reach different lengths in any species. Here, we directly compare the transcriptomes of homologous growth cartilages of the mouse (*Mus musculus*) and bipedal jerboa (*Jaculus jaculus*), which has extremely long metatarsals of the feet and ‘mouse-like’ arms. When we intersected gene expression differences in metatarsals of the two species with expression differences in forearms, we found that about 10% of all orthologous genes are associated with disproportionate elongation of jerboa feet. Among these, *Shox2*, has gained expression in jerboa metatarsals where it is not expressed in other vertebrates that have been assessed. This transcription factor is necessary for proximal limb elongation, and we show that it is sufficient to increase mouse distal limb length. Unexpectedly, we also found evidence that jerboa foot elongation occurs in part by releasing latent growth potential that is repressed in mouse feet. In jerboa metatarsals, we observed higher expression of *Crabp1*, an antagonist of growth inhibitory retinoic acid, lower expression of *Gdf10*, an inhibitory TGFβ ligand, and lower expression of *Mab21L2*, a BMP signaling inhibitor that we show is sufficient to reduce limb bone elongation. By intersecting our data with prior expression analyses in other systems, we identify mechanisms that may both establish limb proportion during development and diversify proportion during evolution. The genes we identified here therefore provide a framework to understand the modular genetic control of skeletal growth and the remarkable malleability of vertebrate limb proportion.

## Introduction

Diversification of limb skeletal proportion from the most primitive four-legged ancestor allowed animals to fly high above land, travel quickly across its surface, and diligently burrow beneath. However, loss-of-function mutations in many genes, including endocrine and paracrine signals and components of intracellular pathways, produce proportionately dwarfed skeletons that suggest a common ‘toolkit’ is required for growth of all long bones^1,2^. We therefore do not yet know which genes are responsible for differential growth, let alone how their expression is regulated to allow for modular development and evolution of the forelimb, hindlimb, and of more than a dozen long bones within each limb.

Quantitative trait loci (QTL) analyses and genome wide association studies (GWAS) can identify genes associated with the variance of limb proportion in a population^3–6^. However, the variance in length of a particular bone does not compare to the scale of length differences among different bones. Bone lengths in a mouse or human, for example, span an order of magnitude from the short toe bones to the long tibia, but here it is difficult to distinguish genes that might be responsible for differential growth from their association with positional identity (e.g. the roles of different *Hox* genes to specify the foot and lower leg). An alternative approach would be to directly compare homologous skeletal elements in two or more species with dramatically different proportions. However, large phylogenetic distances can make it difficult to identify phenotypically-relevant orthologous gene expression differences among extensive expression level and sequence divergence.

We addressed these challenges using the uniquely suitable lesser Egyptian jerboa, *Jaculus jaculus*, and the laboratory mouse, *Mus musculus*, which diverged from a common ancestor ~50 million years ago^7^. Although the jerboa has since acquired extraordinarily elongated hindlimbs with disproportionately long feet, it retained ‘mouse-like’ forelimbs that are a valuable control for divergence unrelated to the evolution of skeletal proportion^8^. We leveraged these morphological similarities and differences, and the relatively close phylogenetic relationship of these two rodents, to directly compare the transcriptomes of homologous growth cartilages (radius/ulna of the forearm and metatarsals of the hindfeet). We then intersected these data to identify gene expression differences that are specifically associated with exaggerated metatarsal elongation in the jerboa.

Although a majority of the growth cartilage transcriptome has diverged since the last common ancestor of mice and jerboas, we found that most gene expression has changed to the same extent in both metatarsals and radius/ulna. Expression of only about 10% of genes has evolved in feet independent of forearms. In addition to genes that are expected to directly increase the rate of limb bone elongation (e.g. *Shox2*), we found differentially expressed genes that suggest latent growth potential in mouse metatarsals is de-repressed in jerboa metatarsals (*Crabp1, Gdf10*, and *Mab21L2*). Our data demonstrate the power of an inter-species transcriptomic approach to directly compare similar and extremely divergent structures in order to identify differences with relevance to a particular phenotype in the vast background of gene expression divergence over macro-evolutionary timescales.

## Results and Discussion

### Interspecies expression differences between metatarsals and between radius/ulna

In order to directly quantify gene expression differences between homologous skeletal elements of the jerboa and mouse, we first annotated a set of 17,464 orthologous genes comprised of at least one exon with no frameshift, missense, or nonsense amino acid differences using CESAR^9^ and the *M. musculus* (mm10) and *J. jaculus* (JacJac1.0) genome assemblies (NCBI). This 1:1 orthologous gene set represents 92.5% of predicted protein coding genes in the jerboa and 79.8% of annotated mouse genes, with the difference likely due to a difference in completeness of the mouse versus jerboa genome assemblies and annotations.

We then isolated mRNA from the distal metatarsal (MT) and distal radius/ulna (RU), including growth cartilage and surrounding perichondrial membrane, from mice and jerboas at postnatal day five (P5), soon after perinatal growth rate differences are established and before senescence is initiated in mouse metatarsals^10^. We determined that jerboa metatarsals elongate about twice as fast as mouse metatarsals from birth (P0) to P5, while the radius/ulna elongate at a more similar rate in both species (Fig. 1). We included both the growth cartilage and perichondrium since crosstalk between the two tissues is critical to control endochondral skeletal elongation^11^.

**Figure 1.**
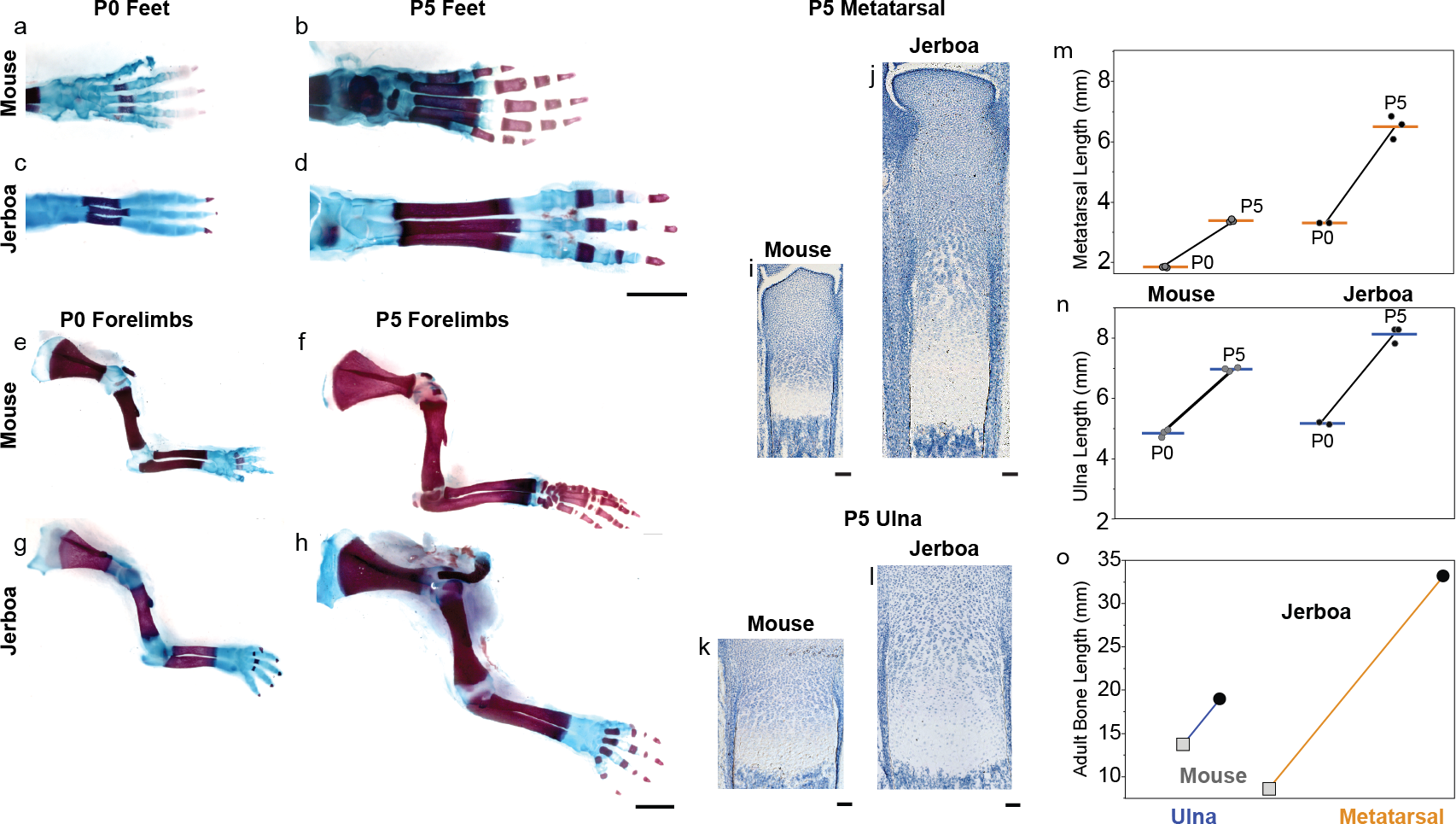
Jerboa metatarsals elongate disproportionately faster during neonatal development. (**a**-**h**) Skeletal preparations of jerboa and mouse limbs. Postnatal day 0 (P0) and 5 (P5) skeletons of mouse (**a**,**b**) and jerboa feet (**c**,**d**). P0 and P5 skeletons of mouse (**e**,**f**) and jerboa arms (**g**,**h**). (**i**-**l**) Histological sections of P5 distal metatarsal (**i**,**j**) and ulna (**k**,**l**) growth cartilages in mouse (**i**,**k**) and jerboa (**j**,**l**). (**m**-**n**) Growth trajectory of neonatal metatarsals (**m**) and ulna (**n**) of mouse and jerboa show that the third (middle) jerboa metatarsal elongates 2.1-times faster and the ulna elongates 1.4-times faster than mouse from P0 to P5. (**o**) Differences in the length of ulna (blue line) and metatarsal (orange line) in adult mouse (squares) and jerboa (circles). Scale bars, **a**-**h** = 2 mm, **i**-**k** = 100 um.

For each of the four sample sets, we sequenced five biological replicates and mapped reads to their respective genome using annotations from our 1:1 orthologous gene set. We analysed differential expression by DESeq2^12^ with an additional custom normalization for gene length differences between species. We then applied a principle components analysis and found that all samples are primarily separated by species (PC1) and secondarily by growth cartilage type (PC2) (Fig. 2a). To reduce the potential effects of technical variation^13,14^ and to reserve samples for independent validation below, we selected for our primary analysis three of the five samples from each dataset that group most closely.

**Figure 2.**
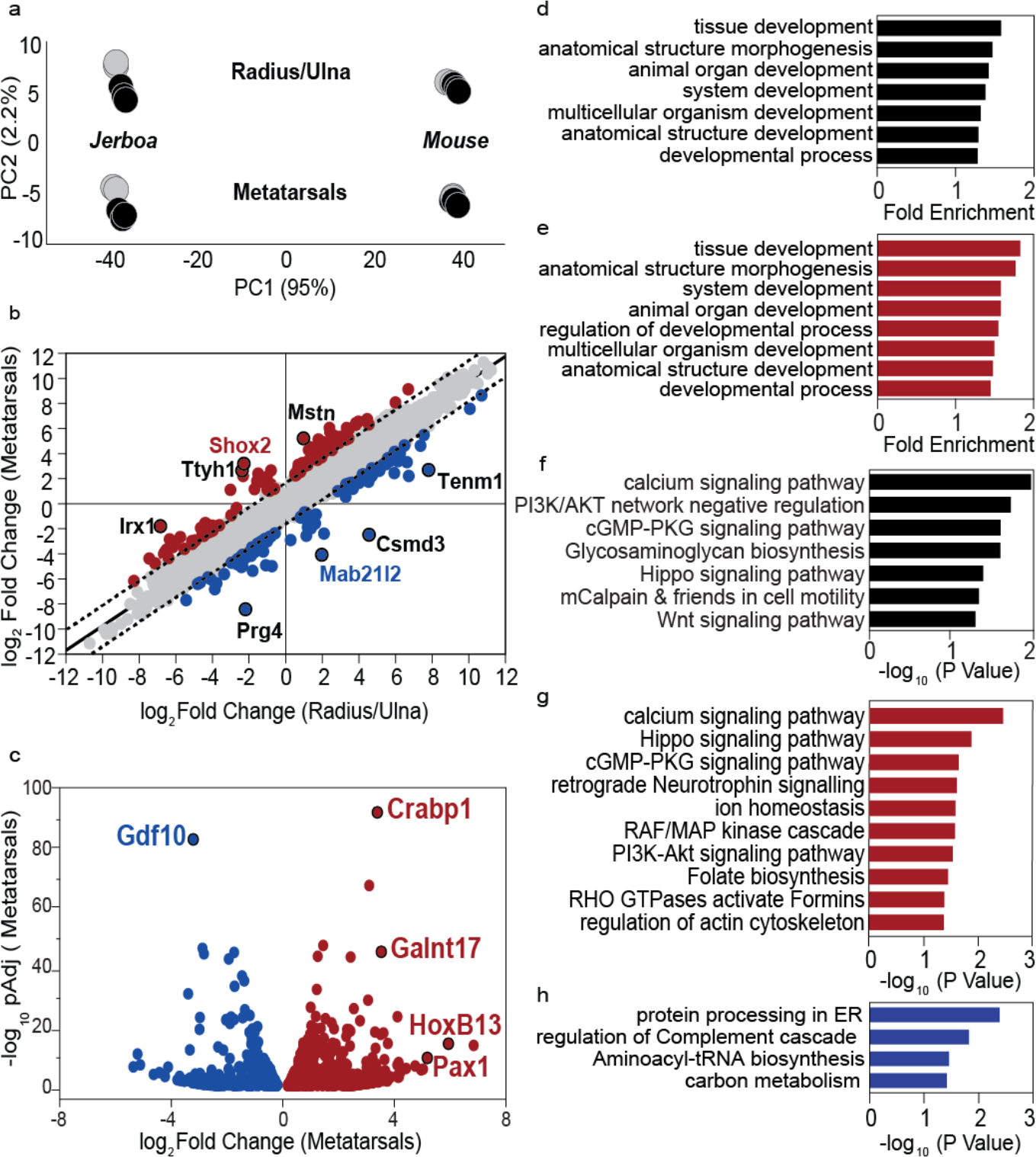
Gene expression profiling identifies 10% of orthologous genes that are associated with disproportionate jerboa metatarsal elongation. (**a**) Principle components analysis (PCA) shows that PC1 (species) explains 95% of the variance and PC2 (tissue type) explains 2.2% of the variance (n=5 each). Black circles denote the three samples chosen for primary analyses. (**b**) Plot of 8,734 genes that are differentially expressed between both metatarsals (MT) and radius/ulna (RU) of jerboa compared to mouse. Of these, 8,493 are equivalently different between species in the MT and RU (grey points) and lie within the 99% confidence interval (dashed line) of the linear fit (solid black line, slope=0.977). The 241 genes outside of the confidence interval show non-equivalent expression differences in the MT and RU. Eight genes with least-correlated differences between MT and RU are labeled. (**c**) 1,514 genes that are significantly differentially expressed between jerboa and mouse MT and not RU. Genes that are higher in jerboa MT than mouse MT are in red and lower are in blue in (**b**) and (**c**). (**d**-**e**) A selection of biological processes GO terms that are significantly enriched (padj <0.05) among (**d**) all 1,755 genes that are associated with disproportionate MT elongation and (**e**) 969 genes that are higher in jerboa MT. (**f**-**h**) Biological signaling pathways that are significantly enriched (padj <0.05) among (**f**) all 1,755 genes, (**g**) 969 genes that are higher, and (**h**) and 786 that are lower in jerboa MT than mouse MT.

As expected given the evolutionary divergence, we indeed found that most genes are differentially expressed between jerboa and mouse metatarsals and/or radius/ulna [58.6% and 57.6% of transcripts respectively with an adjusted p-value (padj) <0.05] (Supplementary Table 2). Of these, however, a majority (~83%) are equivalently different between species in both skeletal elements (Fig. 2b; m=0.977; b=0.025; R^2^=0.930; Supplementary Table 3). Gene expression in these two growth cartilages of the limb skeleton is therefore highly coordinated, even after ~50 million years of divergence since the last common ancestor and regardless of exaggerated jerboa metatarsal elongation. This suggests that many gene regulatory mechanisms are common to multiple cartilages, and mutations that altered DNA binding sites or expression of transcription factors also altered target gene expression coordinately in metatarsals and radius/ulna. These results illustrate the value of directly comparing divergent and more conserved structures of the same tissue composition, because genes that differ equivalently in both locations cannot explain the disproportionate rate of jerboa metatarsal elongation.

Exclusion of these genes revealed a much smaller set (about 10% of all orthologous genes) that likely contribute to the evolution of skeletal proportion, since expression differences between the two species do not correlate in these two skeletal elements. These include 241 genes with non-equivalent expression differences in both the metatarsals and radius/ulna that could contribute to disproportionate growth rate differences between species at both locations (Fig. 2b and Supplementary Table 3). An additional 1,514 genes differ significantly between the metatarsals of the two species but not between the radius/ulna (Fig. 2c and Supplementary Table 4). These include genes that are not detected in the radius/ulna of either species as well as genes with expression that seems constrained in these forearm elements and divergent in metatarsals (Supplementary Fig. 1). Thus, in total, we identified 1,755 genes that are associated with the disproportionate acceleration of jerboa metatarsal growth, which provide the foundation for a mechanistic understanding of the modular nature of skeletal proportion.

Traditionally, differential microarray and RNA-Seq analyses have been validated by quantitative reverse transcriptase PCR (qRT-PCR) of a subset of genes^15,16^. However, qRT-PCR is unsuitable for a direct comparison between species due to the potential for differences in primer annealing affinities and possible differences in the expression of reference transcripts used for normalization^17^. We therefore chose instead to repeat DESeq2 using the two independently collected samples for each growth cartilage that were not included in the primary analysis (Supplementary Tables 5-8) and to perform RNAScope *in situ* detection of transcript expression detailed below. Despite the variance between these and the primary samples, there is a strong correlation among the genes that are associated with the independent acceleration of jerboa metatarsal growth, including all genes we discuss in detail below (p=1.7E-106, Fisher’s exact test; Supplementary Fig. 2).

### Functional Enrichment Among Genes Associated with Differential Growth

In order to broadly assess the classifications of gene functions associated with accelerated jerboa metatarsal elongation, we queried gene ontology (GO) and biological pathway enrichments. Among all 1,755 genes that are associated with the independent acceleration of jerboa metatarsal growth, we found significantly enriched GO terms for biological processes associated with multicellular development and morphogenesis, consistent with the classification of skeletal maturation and growth as developmental processes (Fig. 2d). When we separately analyzed the set of genes that are higher or lower in jerboa metatarsals compared to mice, we found that most of these terms are enriched among genes with higher expression in jerboa metatarsals (Fig. 2e), and no terms are enriched among those with lower expression than in mice (Supplementary Table 9). When we queried biological pathway enrichment among all 1,755 genes, we found that significant enrichments included molecular pathways that were previously associated with tissue growth and/or skeletal growth, including *Hippo* signaling and *Wnt* signaling (Fig. 2f)^18–20^. Similar to our GO term analysis, many of these were also enriched among genes with higher expression in jerboa metatarsals and not among those with lower expression (Fig. 2g,h). A complete analysis of GO term and biological pathway enrichment can be found in Supplementary Tables 9-12.

### Mechanisms of differential growth acceleration and differential growth repression

The disproportionately exaggerated length of the foot in jerboa compared to mouse may result from a difference in molecular mechanisms that directly accelerate growth rate (e.g., more growth factor expression in jerboa metatarsals compared to mouse metatarsals). It is also possible that there are mechanisms that repress the rate of mouse metatarsal elongation, and these ‘brakes’ may have been released in jerboa metatarsals. Here, we highlight differentially expressed genes that suggest both differential acceleration and repression may explain growth rate differences that establish adult proportion during development and evolution.

Among the 241 genes with non-equivalent expression differences in both forearms and in metatarsals, we focused on genes that are least correlated in that they lie furthest from the linear regression of correlated expression differences in both growth cartilages. These include a key example that is likely to accelerate jerboa metatarsal elongation. *Shox2* is increased 9.1-fold in jerboa metatarsals compared to mouse (padj=2.2E-16) and decreased 4.9-fold in jerboa radius/ulna (padj=1.3E-50; Fig. 2b). Loss-of-function mutation of the *Short stature homeobox* transcription factor, *Shox*, causes short stature in a variety of human genetic disorders, including Turner syndrome^21,22^ and Langer syndrome^23^. Although *Shox* is not present in rodent genomes, its paralogue, *Shox2*, is necessary for elongation of the proximal mouse limb skeleton. It is not, however, required for growth of the hands and feet where its expression has never been detected in multiple vertebrate species^24–26^. To confirm this unusual domain of jerboa *Shox2* expression, we performed RNAScope *in situ* hybridization to detect individual transcripts in sections of growth cartilage from each species^27^. We observed no *Shox2* expression in mouse metatarsals, as expected, but *Shox2* appears in the proliferative zone and perichondrium of jerboa metatarsals as in the radius/ulna of both species (Fig. 3a,b) and Supplementary Fig. 3c,d).

**Figure 3.**
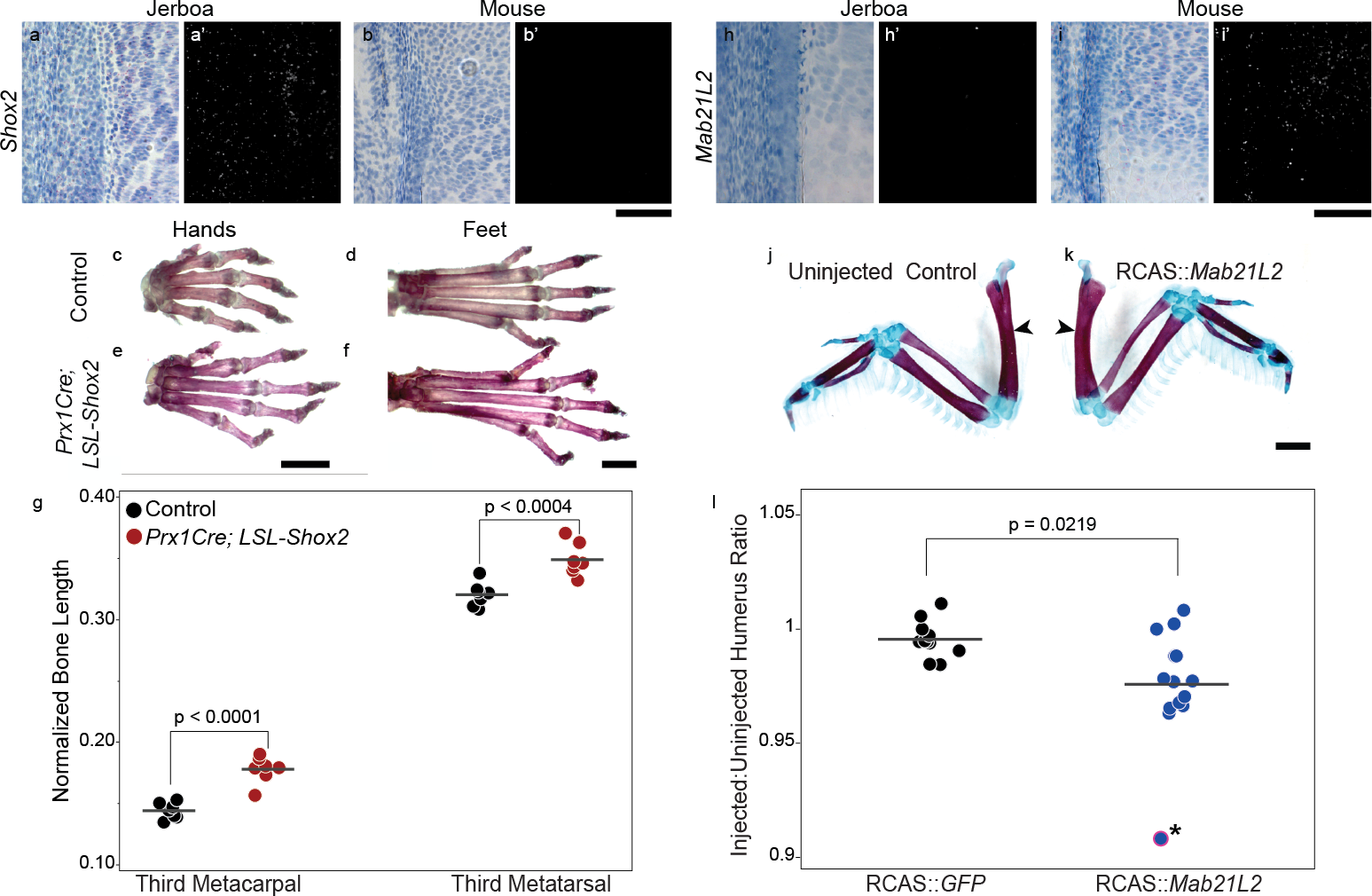
*Shox2* is sufficient to increase and *Mab21L2* is sufficient to reduce limb skeletal elongation. (**a**-**b**) *Shox2* is expressed in jerboa metatarsals where it is not present in mouse. (**c**-**g**) Metacarpal and metatarsal lengths are increased in *Prx1Cre;LSL-Shox2* (*Shox2* overexpressing) mice at eight weeks of age compared to control littermates. In (**g**), measurements are normalized to the skull length of each individual, and p-values are derived from a paired t-test between matched sex littermates. *Shox2* over-expressing metacarpals are 23.6% and metatarsals are 9.1% longer than controls. (**h**-**i**) *Mab21L2* expression is highly reduced in P5 jerboa MT compared to mouse. (**j**-**k**) Skeletal preparations show the left unmanipulated and right RCAS::*Mab21L2* infected wings of a representative chicken embryo. (**l**) Ratio of the lengths of unmanipulated to infected humerus in RCAS::*GFP* control and RCAS::*Mab21L2* chicken embryos shows a 1.9% length reduction in *Mab21L2* overexpressing humerus (p-value derived from a Kolmogorov-Smirnov asymptotic test). The RCAS::*Mab21L2* outlier marked with an asterisk (*) was excluded from statistical analysis. Scale bars, **a**-**b** and **h**-**i**= 50 um, **c**-**f**= 2 mm, **j**-**k**= 200 um.

In order to test the hypothesis that *Shox2* is sufficient to increase the length of distal skeletal elements, we measured metacarpals and metatarsals of mice in which *Shox2* was overexpressed in limbs of mice expressing *Prx1-Cre* and *Rosa26^CAG-loxSTOPlox-Shox2^* transgenes (referred to here as LSL-*Shox2*)^28–30^. We observed a 24% increase in metacarpal length and a 9% increase in metatarsal length (Fig. 3c-g), consistent with observations that the chicken ortholog of *Shox* is also sufficient to increase wing digit length when virally misexpressed in developing chicken wings^31^. Together, these data suggest that the evolutionary co-option of a novel *Shox2* expression domain in jerboa metatarsals may have contributed to their increased elongation rate.

We also discovered evidence of released growth repression in jerboa metatarsals. For example, expression of *Mab21L2* is 16.8-fold decreased in jerboa metatarsals (padj=1.8E-26) and 3.9-fold increased in the radius/ulna (padj=3.9E-16) compared to mice. RNAScope *in situ* hybridization confirmed expression in resting zone and proliferative chondrocytes as well as the perichondrium of mouse metatarsals, whereas *Mab21L2* expression was absent in jerboa metatarsals (Fig. 3h,i). *Mab21L2* interacts with Smad transcription factors to inhibit BMP signaling^32^. BMP signaling promotes bone growth, which is evident from loss-of-function mutations in BMP receptors that cause severe chondrodysplasia in mice^33,34^ and from overexpression of *Bmp2* and *Bmp4* in chicken embryo limbs that each increase the size of skeletal elements^35^. Reduced *Mab21L2* expression in jerboa metatarsals is exciting in light of the fact that its expression is also reduced in tissues associated with the elongated digits of the fetal bat wing compared to the short first wing digit, bat hindlimb digits, and all mouse digits^36^. The authors suggested a role for *Mab21L2* in the establishment of anterior-posterior asymmetry in the bat wing. We predict that *Mab21L2* inhibits elongation of short digits, presumably by repressing BMP signaling, and this growth repression is convergently alleviated in part by decreased *Mab21L2* expression in both the bat wing and jerboa foot. The result in the bat would promote disproportionate elongation of the posterior digits relative to digit I and indeed give the appearance of altering symmetry. However, a specific role for *Mab21L2* in growth cartilage elongation remains unknown, as the gene is also required for embryo viability^37^.

To test the hypothesis that *Mab21L2* expression is sufficient to repress skeletal elongation, we leveraged an approach to misexpress genes by RCAS retroviral infection in developing chicken embryos^38^. When injected into the early developing limb bud, the replication competent RCAS retrovirus broadly infects limb tissues, including the developing skeleton^39^. Since elongation of the right and left limbs is highly coordinated to maintain symmetry during development^40^, any significant deviation from the length of contralateral non-infected limb skeletal elements, compared to control infection, is evidence that a gene is sufficient to promote or inhibit growth. We found that the humerus of limbs that over-express RCAS:*Mab21L2* is on average 1.9% shorter, in contrast to the symmetric limbs of control RCAS:*eGFP* infected chicken embryos (p=0.0219, Kolmogorov-Smirnov asymptotic test; Fig. 3j-l). Consistent with our observations, two presumed gain-of-function missense mutations that appear to stabilize the human MAB21L2 protein each cause shortening of the proximal limb skeleton in addition to eye malformations^41^. Together, these data suggest that *Mab21L2* is indeed sufficient to inhibit skeletal elongation, and decreasing its expression may have contributed to the convergent evolution of an elongated distal skeleton in the jerboa hindlimb and bat forelimb. That Mab21L2 appears to be a relatively weak BMP inhibitor might reflect the fact that there are multiple intracellular effectors of BMP signaling that can compensate for one another^42,43^ contrasting the potent inhibition by Noggin^44,45^ or Gremlin^46,47^, which each block ligand-receptor interactions. The relatively weak inhibitory effect might make *Mab21L2* amenable to processes of natural selection that can favor subtler multigenic changes over time.

There is additional support for the hypothesis that differential growth is achieved in part by differences in growth repression among the 1,514 genes that are differentially expressed between jerboa and mouse metatarsals and not between radius/ulna (Fig. 2c). *Cellular retinoic acid binding protein 1* (*Crabp1*) is expressed 10.6-fold higher in jerboa metatarsals and has the highest statistical significance among this subset of genes associated with the evolution of skeletal proportion (padj=1.5E-92). *Crabp1* has been used as a marker of perichondrium^48–50^, and its expression is indeed stronger in the perichondrium of jerboa metatarsals compared to mouse by mRNA *in situ* hybridization (Fig. 4a,b). However, its function during skeletal development remains unknown in part because no abnormal phenotype was detected in knockout mice^51^, though a precise analysis of bone lengths was not reported.

**Figure 4.**
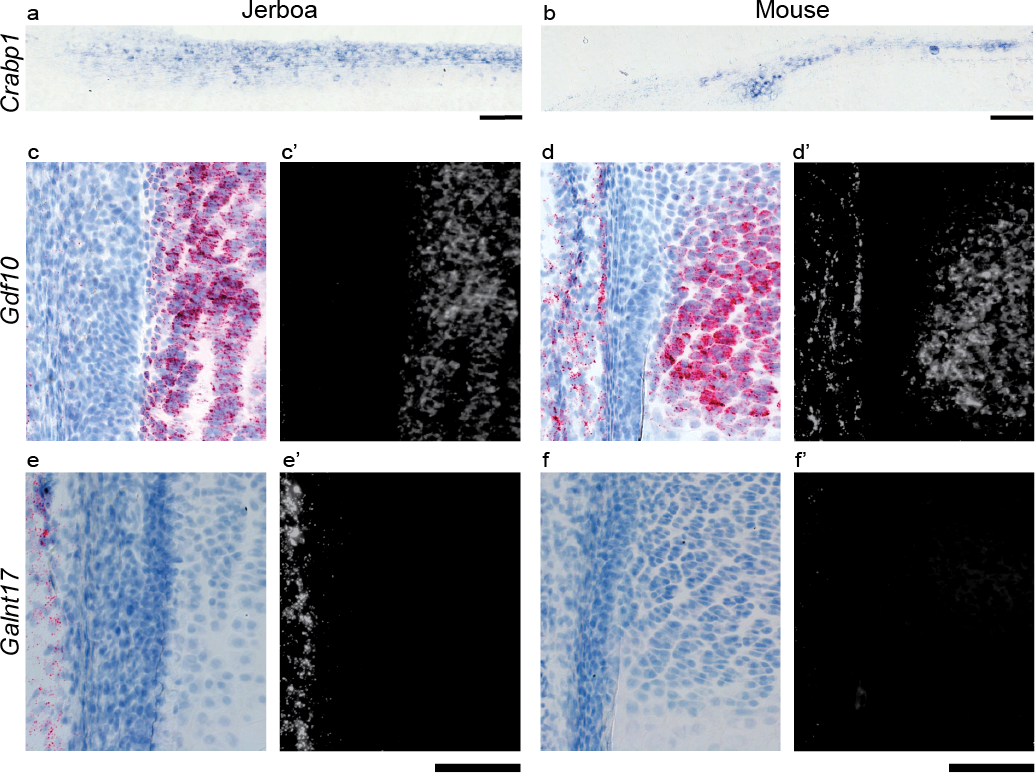
Spatial pattern of *Crabp1, Gdf10*, and *Galnt17* expression in jerboa and mouse metatarsal growth cartilages. Expression patterns in distal growth plates of P5 jerboa (left) and mouse (right). (**a**-**b**) *Crabp1* colorimetric *in situ* hybridization, (**c**-**g**) RNAScope colorimetric and (**c’**-**g’**) fluorescence detection. (**c**-**d**) *Gdf10* and (**e**-**f**) *Galnt17*. Scale bars, **a**-**b**= 100 um, **c**-**h** = 50 um.

In cancer cells, *Crabp1* inhibits the anti-proliferative effects of retinoic acid (RA) by sequestering RA in the cytoplasm^52–54^. Exogenous RA, a potent teratogen, inhibits skeletal elongation in juvenile rats, and pharmacological inhibition of RA signaling accelerates rat metatarsal elongation *in vitro*, suggesting that endogenous RA suppresses metatarsal growth^55^. Intriguingly, *Gdf10/Bmp3b*, an inhibitory TGFβ ligand, is one of two genes upregulated both by RA or an RA Receptor γ agonist in mouse limb culture^56^. We show that *Gdf10* has the highest statistical significance among genes that are lower in jerboa metatarsals compared to mouse (9.3-fold lower, padj=1.3E-83; Fig. 2c). *Gdf10* mRNA is localized to the proliferative zone of both growth cartilages in each species, and strong expression was observed in mouse metatarsals, consistent with our RNA-Seq results (Fig. 4c,d and Supplementary Fig. 3a,b). Together, these data suggest that endogenous RA signaling may repress growth in mouse metatarsals, perhaps in part by increasing *Gdf10* expression. The repressive effect may be alleviated in jerboa metatarsals by increasing *Crabp1* expression to sequester RA and decrease *Gdf10* expression.

Among the genes that differ between metatarsals and not between radius/ulna of the two species, we also found higher expression of two developmental transcription factors in jerboa metatarsals that have not been reported in the distal mouse limb, *Pax1* (62.7-fold higher, padj=6.2E-16) and *HoxB13* (37-fold higher, padj=3.1E-11) (Fig. 2c). In the developing mouse embryo, *Pax1* expression is restricted to the proximal-most limb where it is required for elongation of the acromion, a protrusion of the scapula that articulates with the clavicle, but *Pax1* knockout mice have no other limb abnormality^57^. Similarly, although *HoxB13* is expressed during development and regeneration in amphibian limbs^58^, its expression is not detected in mouse limbs by our analyses or in prior studies^59,60^. Our RNAScope analysis of jerboa tissue sections indeed revealed expression of both genes in connective tissue that lies between adjacent metatarsals of the jerboa foot (Supplementary Fig. 4). For *HoxB13*, this is consistent with previous observations that, in general, *Hox* genes are not expressed within the growth cartilage but rather in adjacent connective tissues and therefore influence growth non-autonomously^61,62^.

We also observed that *Galnt17*, an N-acetylgalactosaminyltransferase among 28 genes deleted in short-statured patients with Williams-Beuren syndrome^63^, is increased 11.7-fold in jerboa metatarsals compared to mouse (padj=2.2E-46). Although *Galnt17*, also known as *Wbscr17*, has no reported growth cartilage function, the International Mouse Phenotyping Consortium annotated a ‘short tibia’ phenotype, suggesting that *Galnt17* might also promote skeletal elongation. Its expression in the jerboa foot also appears in connective tissue that lies adjacent to the perichondrium surrounding each of the three metatarsals (Fig. 4e,f). Together with the expression of *Pax1* and *HoxB13*, this raises the possibility that exaggerated elongation of jerboa metatarsals may be influenced by surrounding connective tissues. Alternatively, these novel expression domains in the jerboa foot may underlie other unknown connective tissue differences between the species.

### Meta-analysis of multiple differential growth datasets

It is not known if similar genetic mechanisms establish limb skeletal proportion during development of an individual, vary proportion in a population, and/or diversify proportion over many millions of years. We therefore tested whether there is a significant overlap between the genes we identified as associated with the evolution of skeletal proportion and three other datasets that are associated with the development of limb proportion in laboratory rodents or the variance of body proportion in a human population. Such correlations might reveal a common set of genes that are recurrently utilized to establish growth rate differences in the vertebrate limb.

First, an analysis of mouse and rat growth plate RNA-Seq identified differentially expressed genes that are associated with the slowing of tibia growth rate as the growth plate matures in juvenile animals of both species^19^. Second, the same study also identified differentially expressed genes associated with growth rate differences by location in the rapidly elongating proximal tibia growth plate of young animals versus the slowly elongating distal phalanx of the same individuals. Third, though not a direct measure of skeletal proportion within limbs, the human sitting height ratio (SHR) GWAS identified SNPs associated at a population level with leg length as a proportion of total body length^5^.

We first identified genes within each of these datasets that were assigned the same gene name in our 1:1 jerboa and mouse orthologous reference set. Of these, there are 287 differentially expressed genes associated with growth rate slowing during tibia maturation, 493 genes that are associated with growth rate differences by location, and 99 unique genes assigned to SNPs with nominal significance in the human sitting height ratio GWAS (Supplementary Table 13). Using our orthologous reference set as background for comparison to genes associated with the evolutionary acceleration of jerboa metatarsal elongation, we found significant correlation with genes associated with tibia maturation as well as genes that are differentially expressed by location (p=5.4E-03 and 3.6E-06, respectively; Fisher’s exact test) but no correlation with sitting height ratio associated genes (p=0.41). We then intersected the three correlating datasets and found thirteen genes that are associated with the evolution of skeletal proportion, slowed growth rate during maturation, and growth rate differences by location (Table 1). Six of these have mouse mutant phenotypes that include bone length alterations, skeletal malformations, bone mineral deficiencies, and/or body size abnormalities that may or may not be a direct result of skeletal growth perturbation. Of particular interest among these, a SNP near the Natriuretic Peptide Receptor 3, *Npr3*, is also associated with increased leg length as a proportion of total body length in humans^5^.

**Table 1:** Genes that are associated with the development and evolution of limb skeletal proportion. Asterisk designates *Npr3*, which also appears in the human SHR GWAS.

| Gene Symbol | Gene Name | Known Skeletal/Body Size Phenotypes |
| --- | --- | --- |
| *Alpl* | alkaline phosphatase, liver/bone/kidney | Abnormal long bone metaphysis morphology<sup>64</sup><br>Decreased long bone epiphyseal plate size<sup>64</sup><br>Decreased bone mineral density<sup>65</sup><br>Increased bone resorption<sup>66</sup><br>Decreased body size<sup>67</sup> |
| *B2m* | beta-2 microglobulin | None reported |
| *Baiap2l2* | BAI1-associated protein 2-like 2 | None reported |
| *Enpp2* | ectonucleotide pyrophosphatase/phosphodiesterase 2 | None reported |
| *Ibsp* | integrin binding sialoprotein | Abnormal trabecular bone morphology<sup>68</sup><br>Decreased bone mineral density<sup>68</sup><br>Increased bone resorption<sup>69</sup><br>Decreased body size<sup>68</sup><br>Short femur<sup>68</sup> |
| *Lmcd1* | LIM and cysteine-rich domains 1 | None reported |
| *Mcoln3* | mucolipin 3 | Decreased body size<sup>70</sup> |
| *Myh10* | myosin, heavy polypeptide 10, non-muscle | Abnormal body size<sup>71</sup> |
| *Nlgn3* | neuroligin 3 | None reported |
| *Npr3\** | natriuretic peptide receptor 3 | Increased width of hypertrophic chondrocyte zone<sup>72</sup><br>Delayed endochondral bone ossification<sup>72,73</sup><br>Increased length of long bones<sup>72,74</sup><br>Elongated metatarsal bones<sup>74</sup><br>Increased body<sup>72,74,75</sup> |
| *Nprl3* | nitrogen permease regulator-like 3 | None reported |
| *Tbx18* | T-box18 | Abnormal skeleton morphology<sup>76,77</sup><br>Decreased body length<sup>77,78</sup> |
| *Ttll9* | tubulin tyrosine ligase-like family, member 9 | None reported |

## Conclusion

The limb skeleton is highly modular with dozens of individual long bones elongating at different rates during development, and each rate has been individually tuned to diversify skeletal proportion across species. The high concordance between genes associated with the development and evolution of skeletal proportion suggests that many of the same genes that establish proportion may also diversify proportion. Among these, some are familiar in the context of skeletal growth while many others, such as *Mab21L2*, had no prior known growth plate function. Our work therefore demonstrates the value of an unbiased approach to prioritize functional analyses of previously understudied genes with potential to reveal new mechanisms of skeletal growth control. Additionally, our identification of genes that have novel expression domains in jerboa metatarsals, such as *Shox2*, suggests that some genes have acquired an ability to accelerate growth rate in locations where they are not typically expressed in other species. All together, these are an ensemble of genes whose modular expression within the skeleton has the potential to constrain growth in some bones while tuning the growth of others to bring about the extraordinary malleability of limb form and function.

## Acknowledgements

We are grateful to Dr. Wayne Pfeiffer and Dr. Mahidhar Tatineni at the San Diego Super Computing Cluster (SDSC) for computational assistance and to Dr. David Traver for microinjector usage. We are grateful to Dr. Fan Wang (Duke University) for supplying the *Rosa26*^*CAG-loxSTOPlox-Shox2*^ mice. We thank Dr. Stephen Hughes and Andrea Ferris (NCI-NIH) for RCAS vectors. We also thank Drs. C. Tabin, J. Posakony, and J. Monda for comments on the manuscript. This work used the Extreme Science and Engineering Discovery Environment (XSEDE) at SDSC, which is supported by National Science Foundation grant number ACI-1548562. This work was also supported by Natural Sciences and Engineering Research Council grant RGPIN/355731-2013 to JC and by a Searle Scholar Award from the Kinship Foundation, a Pew Biomedical Scholar Award from the Pew Charitable Trusts, a Packard Fellowship in Science and Engineering from the David and Lucile Packard Foundation, and the National Institutes of Health under award number R01AR075415 awarded to KLC.

## Methods

### Animals

Jerboas were housed and bred as previously described^79^. CD-1 mice and fertilized chicken eggs were obtained from Charles River Laboratories (MA, USA). All animal care and use protocols were approved by the Institutional Animal Care and Use Committee (IACUC) of the University of California San Diego or the Life and Environmental Sciences Animal Care Committee of the University of Calgary.

### Skeletal preparations and measurements

Freshly dissected chicken wings were fixed while rocking a room temperature in 100% ethanol for 24 hours followed by 100% acetone for 24 hours. Specimens were then stained for 3-5 days with alcian blue/alizarin red staining solution (one volume each of 0.3% alcian blue-8GX in 70% ethanol, 0.1% alizarin red-S in 95% ethanol, and glacial acetic acid plus 17 volumes of 70% ethanol). After staining, specimens were rinsed with de-ionised water and destained over several days in a graded series of 1% KOH followed by 20%, 50%, and 80% glycerol diluted with 1%KOH until stain was sufficiently cleared from the soft tissues. Specimens were stored and imaged in 100% glycerol at room temperature. Each specimen was then blinded to measure skeletal elements using digital calipers. The length of each skeletal element in each specimen is represented as the average of three independent blind measurements to reduce random error and improve precision.

Mice that were hemizgygous for *Rosa26^CAG-loxSTOPlox-Shox2^* and *Prx1-Cre* were generated as previously described^30^. Controls for this cross were siblings that were hemizygous for only one or neither of these transgenes. Eight-week old mouse skeletons were stained with Alizarin red as follows. Mice were skinned and eviscerated, and internal organs were removed. Mice were fixed in 100% ethanol for 4 days followed by acetone for 3 days rocking at room temperature. The specimens were then incubated in 2% KOH for 5 days to dissolve tissue. The skeletons were subsequently stained with 0.005% Alizarin red (Sigma- Aldrich A5533) in 2% KOH for 3 days until the bones appeared red. Following staining, the skeletons were destained with 20% glycerol in 1% KOH to remove excess stain followed by 100% glycerol. All specimens were blinded for measurements as described above.

### RNA sequencing and analysis

Distal metatarsal (MT) and radius/ulna (RU) growth cartilages with intact perichondrium were dissected in ice cold PBS from freshly euthanized mice and jerboas at postnatal day five. Dissected growth cartilages were treated for three minutes at room temperature in a 25% dilution of Proteinase-K (Qiagen) in PBS to remove muscle and connective tissue. Samples were equilibrated in RNA*later* solution (Qiagen) overnight at 4°C and stored at −80°C until ready for RNA extraction. To extract RNA, growth cartilages were pulverized in a liquid nitrogen cooled chamber. Pulverized samples were further homogenized with QIAshredder columns (Qiagen). Homogenate was treated with Proteinase-K (Qiagen) at 55°C for 10 min. RNA was extracted using the RNeasy Micro kit (Qiagen) following the manufacturer’s protocol that included an on-column DNase treatment step. cDNA libraries were prepared using Illumina TruSeq^®^ stranded mRNA library preparation kit following the manufacturer’s protocol. We multiplexed libraries and sequenced single-end 50-bp reads (SR50) on an Illumina HiSeq-4000 platform at the Institute for Genomic Medicine (IGM) at UC San Diego. Low quality reads and residual adaptor sequences were trimmed using Trimmomatic^80^ (version 0.35).

Using the Coding Exon-Structure Aware Realigner (CESAR)^9^ we aligned exons from jerboa (JacJac1.0, NCBI) and mouse (mm10, NCBI) genomes to generate a custom 1:1 orthologous transcript annotation set with no paralogs. Briefly, we first generated a whole genome alignment between the mouse and jerboa genome assemblies^81^. To filter out exonic alignments that represent potential paralogs or even intact exons from processed pseudogenes, we removed all such alignments using the procedure described in Sharma et al^82^. CESAR was then applied to the filtered genome alignment to annotate mouse exons (Ensembl version 87, longest isoform per gene) in the jerboa genome. Exonic regions that appear only once in the resulting jerboa annotation file were retained. Exons corresponding to a particular mouse transcript were combined together to create the orthologous jerboa gene structure. Lastly, those jerboa genes where all the constitutive exons do not come from the same scaffold and strand were discarded. The resulting orthologous reference gene set contains 17,464 transcripts in both genomes with at least one exon free of frame-shift, mis-sense and non-sense mutations. We mapped MT and RU RNASeq reads to jerboa or mouse genomes using these 1:1 orthology GTF annotations and computed gene counts for each library with the STAR aligner^83^. We were able to uniquely map 78.3-88.4% of mouse and 88.2-92.9% of jerboa SR50 reads to their respective genomes with this approach.

STAR gene counts were used to perform differential expression analysis between jerboa and mouse metatarsals and radius/ulna with DESeq2, a robust approach that employs a negative binomial generalized linear model to identify differentially expressed genes^12^. Since orthologous gene/transcript lengths could vary between jerboa and mouse genomes, we implemented an additional length normalization step in the DESeq2 pipeline to avoid false comparative quantifications resulting from species-specific gene/transcript length variation. To do this, we first calculated gene lengths in number of basepairs in non-overlapping exons of each gene from our 1:1 orthologous GTF annotations. We then created a matrix of these lengths for each gene in each sample and input these into the DESeq2 DESeqDataSet object so that they are included in the normalization for downstream analysis. Principle components analysis (PCA) was performed with default DESeq2 settings to identify variance components associated with our MT and RU comparisons (n=5 in each species). We considered all differentially expressed genes with an adjusted p-value < 0.05 to be statistically significant in our analyses. Hierarchical clustering in Supplementary Figure 1 was performed using DESeq2 rlog-transformed gene counts to stabilize variance. Gene counts for all 17,464 transcripts were considered for the rlog transformation.

### GO term and biological pathway enrichment analysis

Gene Ontology (GO) term enrichment was analyzed using over-representation Fisher’s test with Bonferroni correction in PANTHER^84,85^. Enrichment of biological signaling pathways was assessed using DAVID functional annotation for pathway enrichment^84,86^. For each analysis, we input the 1:1 jerboa-mouse orthology annotation set as a reference/background list.

### Test for significance of overlap between gene lists

Fisher’s exact test was performed using the GeneOverlap package^87^ (version 1.20.0) in R to test the statistical significance of overlap between DESeq2 differentially expressed genes in mouse and jerboa for the three primary samples and for the two independently analyzed samples (Supplementary Figure 2b-c). We also used Fisher’s exact test to evaluate the significance of overlap between gene lists from multiple differential growth datasets in our meta-analysis (Table1).

### *In situ* hybridization of P5 growth-plate sections

Metatarsal (MT) and radius/ulna (RU) growth plate cartilages were fixed in 4% PFA in PBS for 24 hours at 4°C. Samples were allowed to equilibrate in 20% sucrose-PBS for 2 days at 4°C. Cartilages were mounted in Tissue-Tek OCT (Sakura) or SCEM compound (SECTION-LAB Co. Ltd., Japan) and cryo-sectioned at 20-30 um thickness. Species-specific digoxigenin (DIG) labeled riboprobes were generated by *in vitro* transcription of *Crabp1* fragments cloned in pGEMT-Easy (Promega) vector. Colorimetric *in situs* were performed on neonatal limb sections following a standard protocol as previously described^88^. For RNAScope hybridization (Advanced Cell Diagnostics, Inc.), custom designed ZZ probes were used to detect *Shox2*, *Mab21L2*, *Gdf10*, *Wbscr17*, Pax1, and *HoxB13* transcripts in P5 growth cartilages of jerboas and mice. Hybridization was performed following manufacturer’s protocol with RNAscope^®^ 2.5 high definition (HD)-RED kit and BaseScope™ detection reagent kit v2-RED (only for jerboa *Mab21L2*). The red chromogen staining was assessed under visible (DIC) and UV (550 nm) light on an Olympus Model BX61 compound microscope with a 20× N.A. 0.75 objective.

### *GFP* and *Mab21L2* RCAS injections in chickens

RCAS vectors that improve the ease of cloning cargo genes, RCASBP(A)-ΔF1 and RCASBP(A)-ΔF1::GFP, were kind gifts from Drs. Andrea L Ferris and Stephen H Hughes (National Cancer Institute, MD, USA). The 1080 bp jerboa *Mab21L2 (XM*_004655859, NCBI) CDS region was synthesised as a gene-block (Integrated DNA Technologies, Inc.) and cloned between *ClaI* and *MluI* restriction sites of RCASBP(A)-ΔF1. DF-1 chicken fibroblast cells (kind gift of Prof. Cliff Tabin, Harvard University, USA) were used to produce virus. Virus production, concentration and pressurized injections into 2-2.5 day old chicken forelimb buds were performed as previously described^89^.

JMP^®^, Version *14.2.0*. (SAS Institute Inc., Cary, NC, 1989-2019) was used to generate all graphs and to perform the Kolmogorov-Smirnov asymptotic test of significant difference between RCAS::GFP and RCAS::Mab21L2-infected chicken humerus.

## Supplementary Figures

**Supplementary Figure 1.**
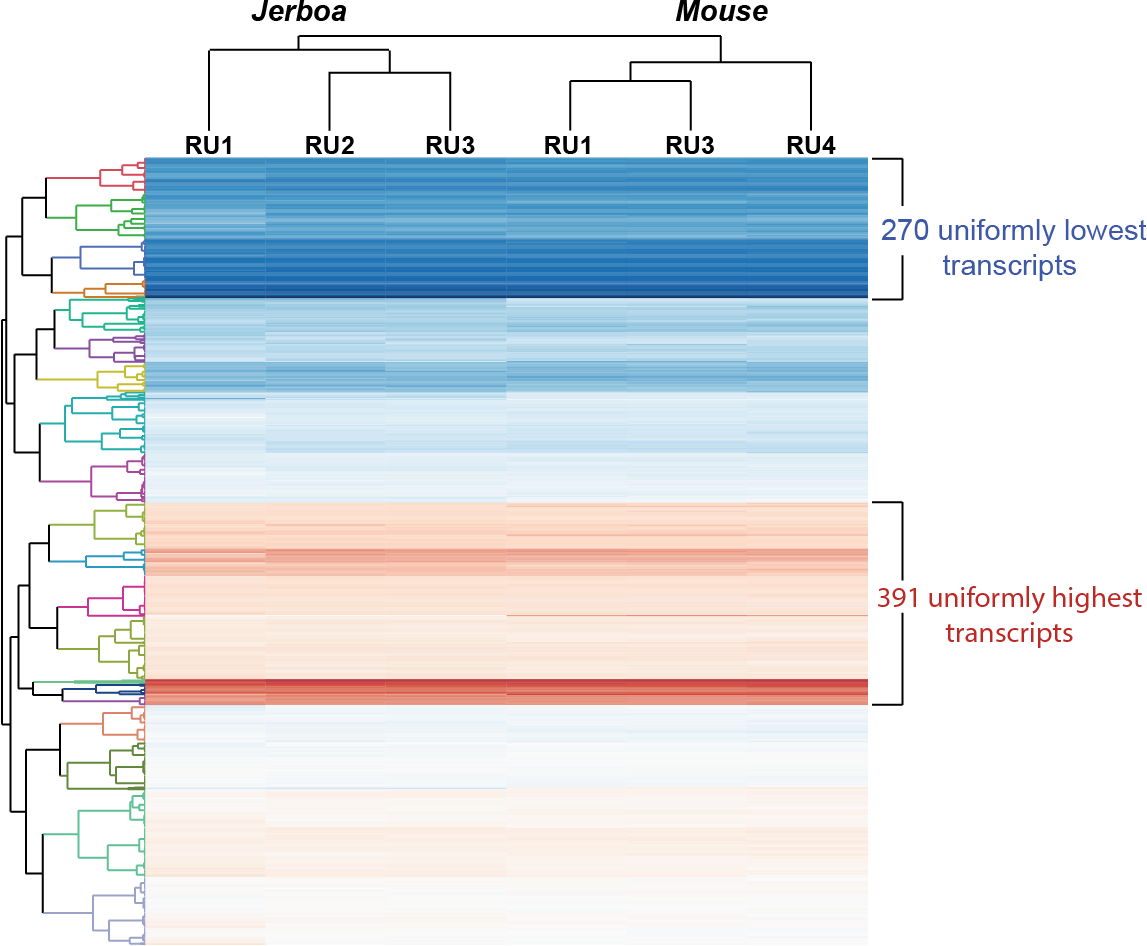
Genes that are differentially expressed between MT but not RU include a cluster with extremely low to undetectable expression in RU and a cluster with high expression in RU. r-log transformed gene counts for three jerboa (left) and mouse (right) RU samples were hierarchically clustered using Ward’s method. Low to high gene expression is marked in blue to red colors respectively. Clusters of genes with lowest or highest expression in RU of the two species are indicated.

**Supplementary Figure 2.**
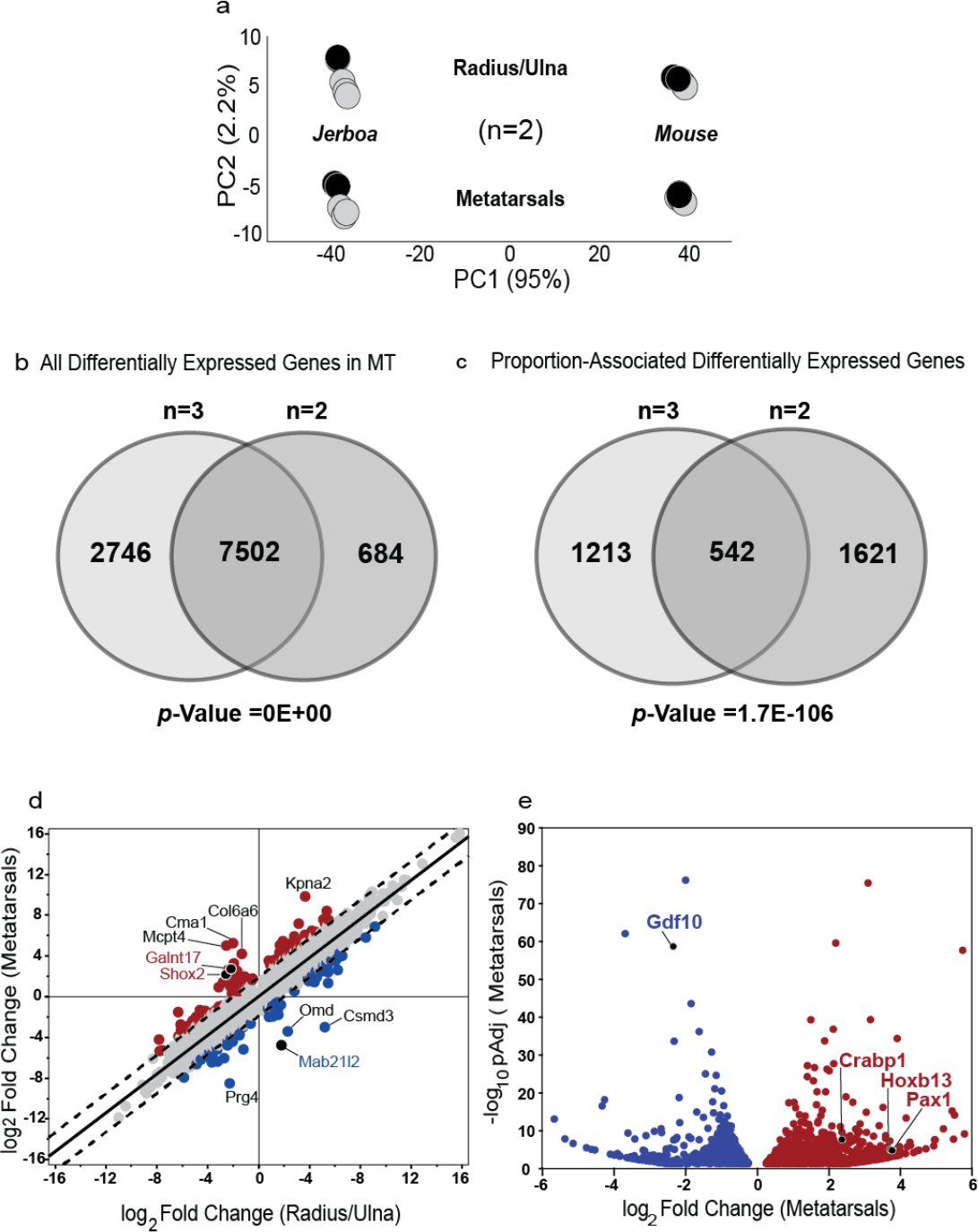
Validation of interspecies RNASeq using two additional independently collected samples. (**a**) Principle components analysis of all five biological replicates with two used for validation marked in black and the three primary samples marked in grey. Venn diagram shows overlap between (**b**) all genes differentially expressed between jerboa and mouse MT and (**c**) the proportion-associated differentially expressed genes (i.e., subtracting genes that are equivalently differentially expressed in MT and RU). Significance of the overlap is indicated and was calculated using Fisher’s exact test. (**d-e**) Plots of gene expression differences in both growth cartilages and differences in MT but not in RU with datapoints highlighted that are discussed in the main text. Slope=0.952 for the correlation between gene expression differences in both MT and RU in (**d**).

**Supplementary Figure 3.**
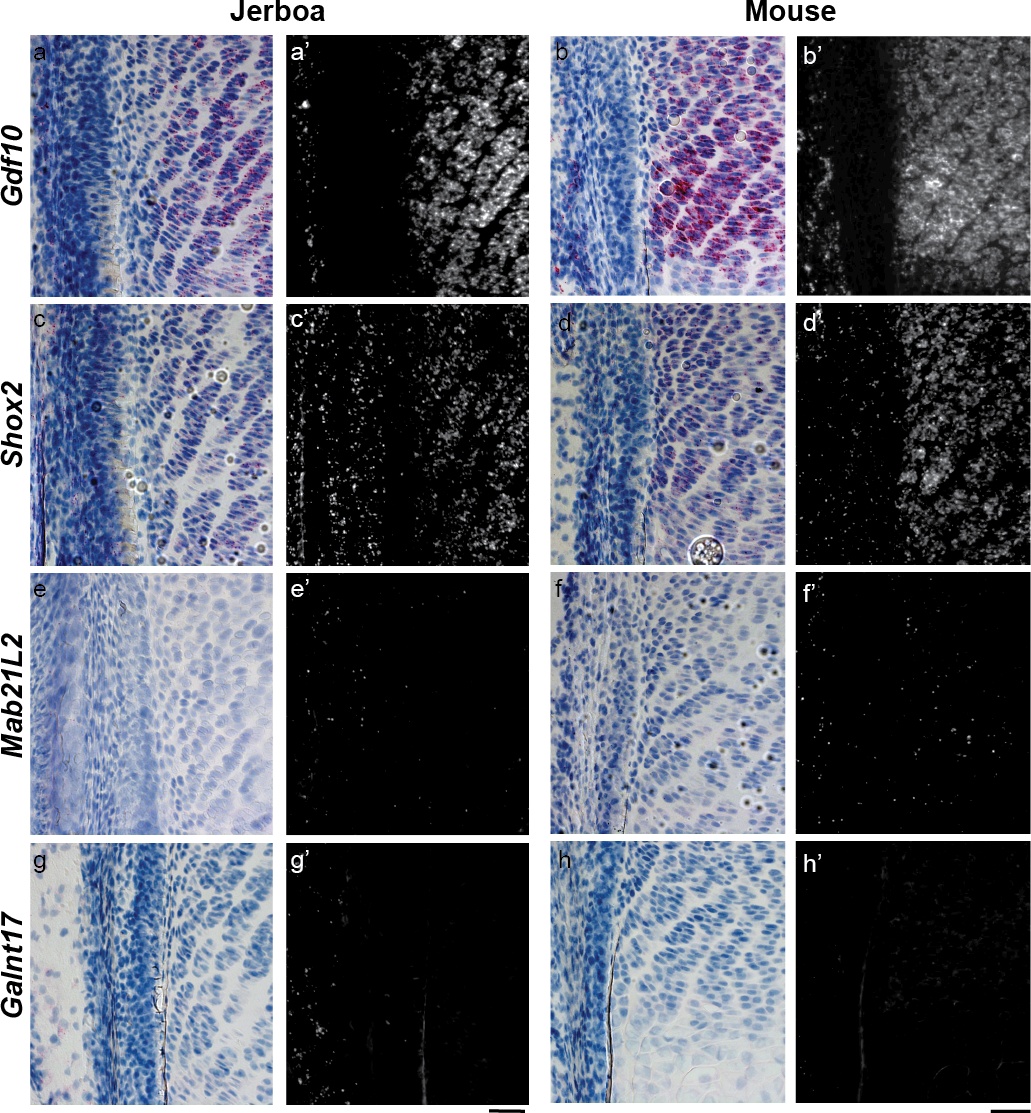
Spatial pattern of *Gdf10, Shox2, Mab21L2*, and *Galnt17* expression in jerboa and mouse radius/ulna growth cartilages. Expression patterns in distal growth plates of P5 jerboa (left) and mouse (right). (**a-h**) RNAScope colorimetric and (**a’-h’**) fluorescence detection. (**a, b**) *Gdf10*, (**c, d**) *Shox2*, (**e, f**) *Mab21L2*, and (**g, h**) *Galnt17*. Scale bars, 50 um.

**Supplementary Figure 4.**
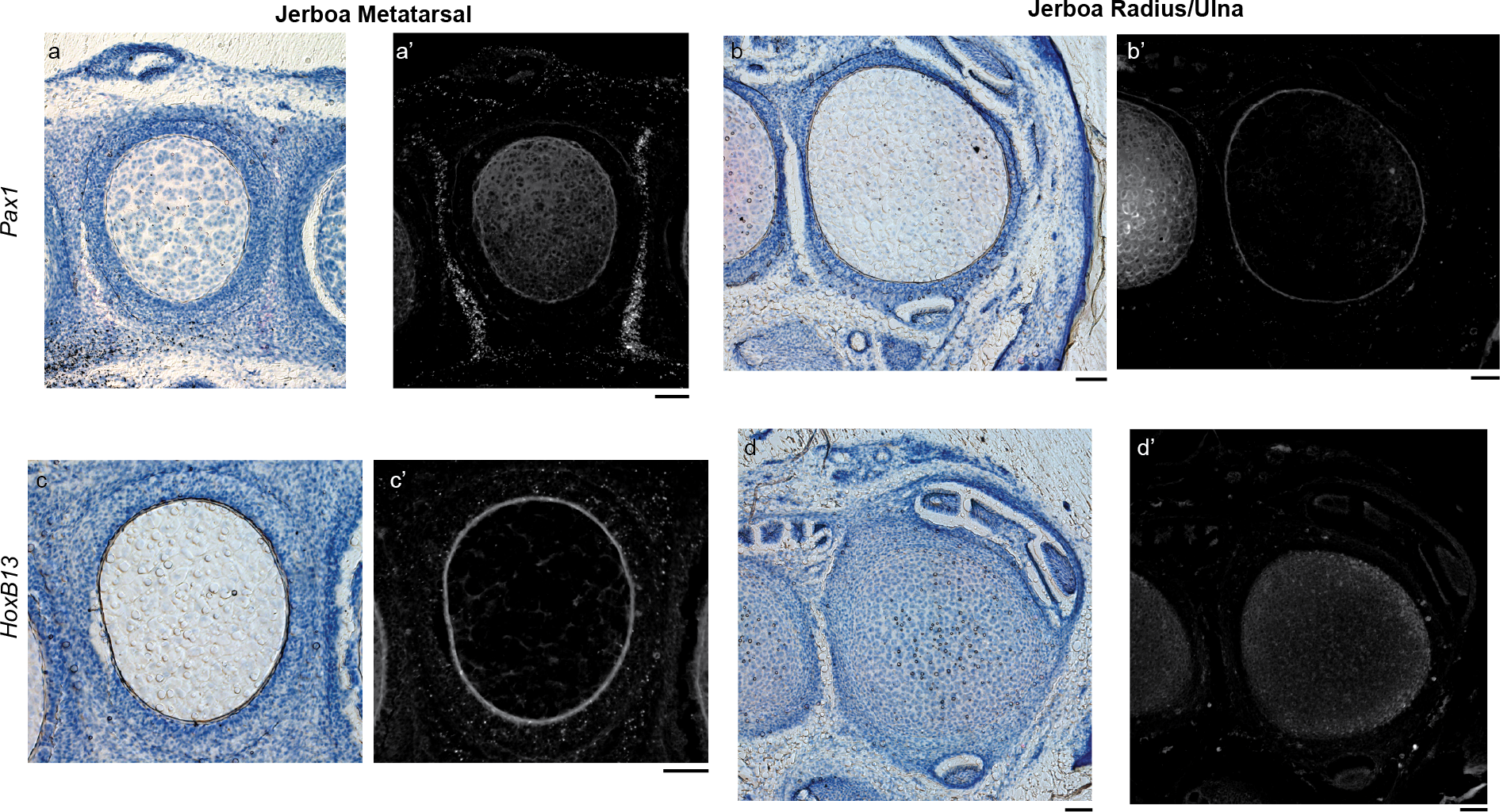
Spatial pattern of *Pax1* and *Hoxb13* mRNA expression in jerboa metatarsal and radius/ulna growth cartilages. Transverse sections through the distal growth cartilage of P5 (**a-b**) metatarsals and (**c-d**) radius/ulna. (**a-d**) RNAScope colorimetric and (**a’-d’**) fluorescence detection shows expression in connective tissue between jerboa metatarsals. *HoxB13* is present as sparse dots throughout the perichondrial connective tissue. Cartilage and/or the edge of cartilage is apparent by autofluorescence as a circle or ring in each.

